# Common ISAV haemagglutinin esterase variants at residues 229 and 230 influence the efficiency of cell-bound receptor destruction and erythrocyte release in Atlantic salmon

**DOI:** 10.64898/2026.06.18.733082

**Authors:** Johanna Hol Fosse, Adriana Magalhães Santos Andresen, Lada Ivanova, Frieda Betty Ploss, Inger Austrheim Heffernan, Simon Chioma Weli, Sonal Jayesh Patel, Petra Elisabeth Petersen, Maria Marjunardóttir Dahl, Debes Hammershaimb Christiansen, Lisa Furnesvik, Knut Falk

## Abstract

Sialic acid-binding viruses often encode receptor-destroying enzymes that modulate infection through host glycan alteration. *Isavirus salaris* (ISAV), the aetiological agent of infectious salmon anaemia, encodes a haemagglutinin esterase with 4-*O*-acetylesterase activity. The ISAV esterase removes the virus-targeted epitope and causes homologous attachment interference, but its broader biological roles remain poorly understood. Several aspects of ISAV receptor engagement and release remain unresolved: Although ISAV agglutinates erythrocytes from both rainbow trout and Atlantic salmon, only rainbow trout erythrocytes elute from this interaction. By contrast, Atlantic salmon erythrocytes remain persistently bound, despite the presence of an active esterase. Here, we used ISAV-erythrocyte interactions to dissect receptor destruction and dissociation across viral genotypes and strains. We uncovered substantial functional diversity in viral permissiveness for Atlantic salmon erythrocyte elution. Common variants of residues 229 and 230 of the haemagglutinin-esterase, located at the distal rim of the P1 esterase pocket, were key determinants of elution permissiveness. Elution further required an active esterase catalytic triad. *In vivo*, Atlantic salmon infected with ISAV carrying the elution-permissive 229N variant showed earlier loss of erythrocyte Neu4,5Ac_2_ than fish infected with an elution-restrictive strain. The loss of Neu4,5Ac_2_ preceded the reduction in ISAV-bound erythrocytes by several days, indicating that receptor removal alone was insufficient to trigger virion release. Targeted sialic acid analyses revealed similar levels of Neu4,5Ac_2_ on Atlantic salmon and rainbow trout erythrocytes. Atlantic salmon erythrocytes additionally expressed di-*O*-acetylated Neu4,5,9Ac_3_. By contrast, brown trout erythrocytes expressed little or no Neu4,5Ac_2_, consistent with their reported inability to support ISAV haemagglutination. Together, our findings give insight into viral and host determinants of ISAV receptor interactions. We demonstrate functional diversity within the ISAV esterase and link virion binding and release to salmonid erythrocyte sialic acid composition. The delay between receptor loss and virion dissociation further supports the existence of additional erythrocyte attachment factors.

## Introduction

Many pathogens use sialic acids as specific attachment receptors. A subset of sialic acid-binding viruses also encodes receptor-destroying enzymes that eliminate virus-targeted host sialic acids. This enzymatic activity limits subsequent virus-receptor interactions at the host cell surface (1).

Infectious salmon anaemia virus (ISAV) is a member of the family *Orthomyxoviridae* (genus *Isavirus*, species *Isavirus salaris*) (2). Its genome comprises eight linear, negative-sense RNA segments, which encode at least ten viral proteins, including the surface glycoprotein haemagglutinin esterase (HE) encoded by segment 6 (3, 4). Receptor binding occurs at the protomer interfaces of the HE homotrimer and requires multimerisation (5). Each HE protomer also contains a sialyl-4-*O*-acetylesterase domain, located approximately 29 Ångström from the receptor-binding site. This esterase specifically hydrolyses 4-*O*-acetylated *N*-acetylneuraminic and *N*-glycolylneuraminic acids (3, 5–7). Pathogenic ISAV variants, defined by heterogeneous deletions in the segment 6 highly polymorphic region (HPR) (8), infect and cause disease in farmed Atlantic salmon (*Salmo salar*). The ISAV esterase mediates global removal of ISAV binding sites from target cell surfaces in infected Atlantic salmon and homologous attachment interference in cell culture (9, 10). Putative additional biological functions of the ISAV esterase remain unresolved. By comparison, the influenza A virus neuraminidase promotes viral fitness by supporting both early and late stages of the infectious cycle, including counteracting respiratory mucus virus inhibition (11–16).

Despite robust ISAV esterase activity both *in vitro* (6, 17) and *in vivo* (9, 10), several reports indicate that ISAV-agglutinated Atlantic salmon erythrocytes fail to elute from the interaction (3). Moreover, in infected Atlantic salmon, ISAV particles remain associated with erythrocytes throughout the active phase of infection, despite extensive loss of ISAV binding sites (9). Two other salmonid species, rainbow trout (*Onchorhynchus mykiss*) and brown trout (*Salmo trutta*), are susceptible to ISAV infection but have not been reported to develop infectious salmon anaemia after natural exposure (18, 19). In contrast to Atlantic salmon erythrocytes, erythrocytes from rainbow trout elute following ISAV agglutination, whereas erythrocytes from brown trout do not support ISAV agglutination (3). The molecular determinants of these species-specific differences have yet to be defined.

Most reports of ISAV receptor-binding and -destroying activities (3, 6, 7, 17, 20) have studied the first-propagated ISAV isolate, Glesvaer/2/90 (21). Prompted by anecdotal observations of occasional elution-permissive isolates, we screened Atlantic salmon erythrocyte elution following ISAV-mediated agglutination across a range of ISAV isolates. We identified elution-permissive strains and analysed HE sequence variation associated with elution phenotypes. Finally, we characterised salmonid erythrocyte *O*-acetylated sialic acid profiles across species and during experimental infection. The findings of our study provide a foundation for understanding host and viral determinants of ISAV-receptor interactions.

## Results

### Atlantic salmon erythrocyte elution following ISAV agglutination varies across isolates

To explore Atlantic salmon erythrocyte elution following ISAV-mediated agglutination, we agglutinated erythrocytes with antigen derived from 13 historical ISAV isolates and screened for elution over a 24-hour period (Table 1, Fig. 1A-B). Consistent with previous reports (3), we observed no erythrocyte elution after agglutination with NO/Glesvaer/2/90 (EU-5) antigen. The same was true for five additional European genotype isolates (EU-1-4, EU-6) (Table 1, Fig. 1B). In contrast, the remaining European genotype isolates (EU-7-10) and all North American genotype isolates (NA-1-3) permitted erythrocyte elution in at least one experiment (Table 1, Fig. 1B). Among elution-permissive isolates, NA antigens supported erythrocyte elution more consistently, with elution observed in 4/5, 4/5, and 5/5 experiments. EU antigens supported elution less consistently, in 1/4, 2/3, 2/4, and 2/5 experiments (Table 1, Fig. 1B). Moreover, NA antigen-agglutinated erythrocytes generally eluted at lower haemagglutination titres than those agglutinated with EU antigen (Fig. 1C). Erythrocytes agglutinated with concentrated virus supernatants, rather than antigen produced in infected cells, displayed elution patterns matching those of corresponding antigens (Fig. 1D), supporting the use of cell-derived antigen as a representative substrate in elution assays.

**Table 1:** Historical ISAV isolates included in elution screen. ^a^Variation from the sequence designated historical EU reference. HE amino acid sequences including residues 17-299 are provided in Table S1. Elution-permissive isolates are shown in red. ^b^HPR typing was performed as recently recommended by Gagné et al. (**22**). ^c^subcultured in RT-gill cells.

| ID | Isolate | HE sequence (17-299)<sup>a</sup><br>[GenBank accession] | Elution<br>(+/total) | HPR<br>type<sup>b</sup> |
| --- | --- | --- | --- | --- |
| EU-1 | NO/Vestvågøy/2163/<br>93 | EU reference<br>[DQ785253.1] | 0/4 | H28.17 |
| EU-2 | NO/Selje/1525/95 | EU reference | 0/4 | H24.21 |
| EU-3 | NO/Hitra/1712/96 | EU Y23H<br>[DQ785245.1] | 0/4 | H19.11 |
| EU-4 | FO/335/01 | EU reference | 0/3 | H14.19 |
| EU-5 | NO/Glesvaer/2/90 | EU I143V T185K<br>[DQ785248.1] | 0/5 | H27.20 |
| EU-6 | NO/Flora/795/00 | EU I29M N201D | 0/4 | H22.19 |
| EU-7 | NO/Solund/288/00 | EU I29M N201D | 1/4 | H22.19 |
| EU-8 | NO/Ørskog/763/01 | EU I29M T185A T230K | 2/3 | H20.11 |
| EU-9 | NO/Midsund/407/99 | EU I29M T185A T230K | 2/5 | H24.24 |
| EU-10 | FO/121/14 | EU I133V T185A T230K L273I A290V<br>[KX823922.1] | 2/4 | H30.17 |
| NA-1 | CA/Canada1SG/97 | NA genotype | 5/5 | H19.7 |
| NA-2 | CA/NBISA01 | NA genotype | 4/5 | H24.17 |
| NA-3 | CA/F679/99<sup>c</sup> | NA genotype | 4/5 | H24.17 |

**Figure 1:**
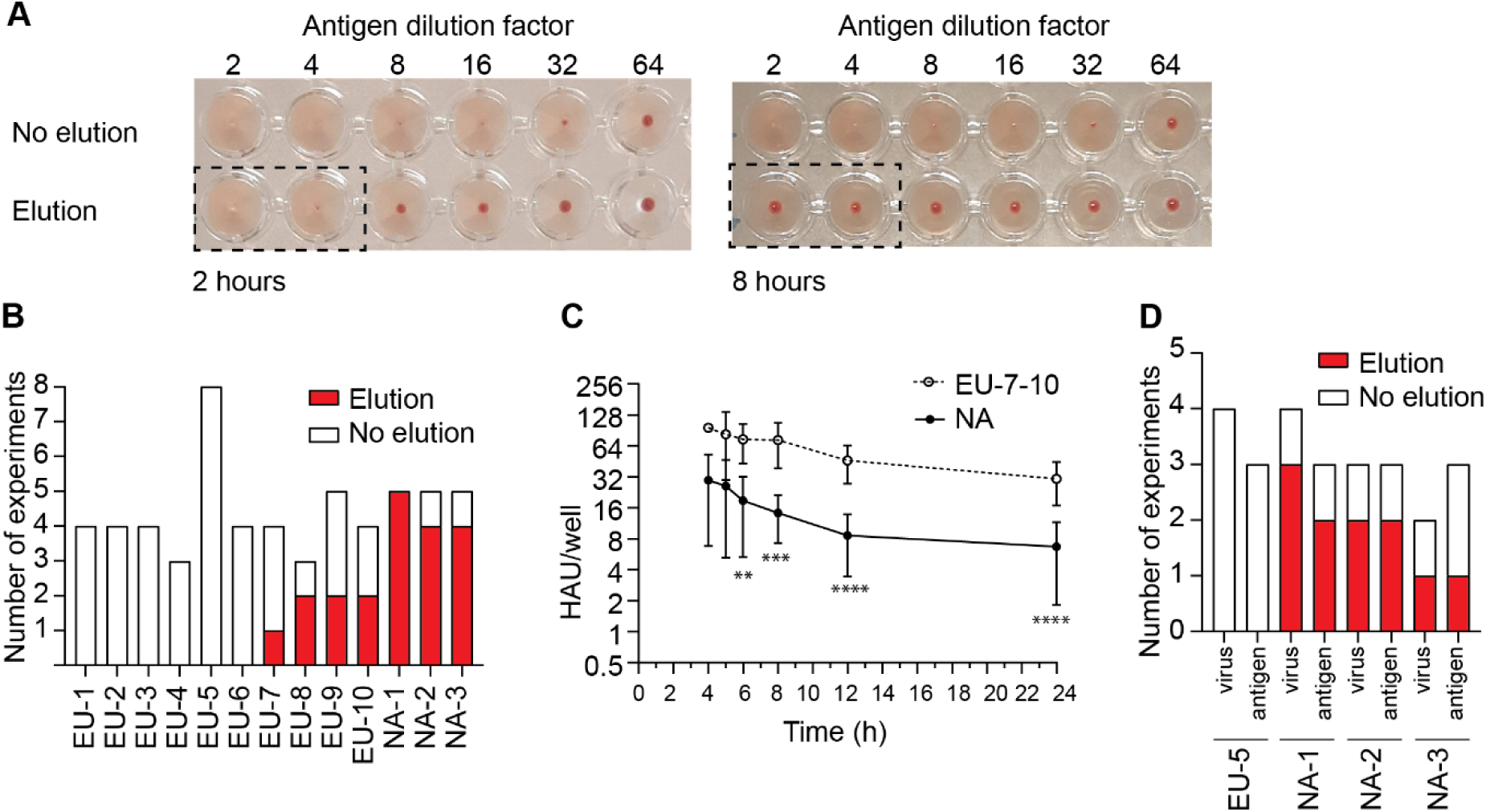
The elution of ISAV-agglutinated Atlantic salmon erythrocytes differs between ISAV subtypes. (A) Representative images from a haemagglutination assay (HA) 2 and 8 hours post-incubation, illustrating typical patterns following agglutination with an isolate that does not permit erythrocyte elution (top) and one that permits elution (bottom, dashed boxes indicate wells with visible elution at 8 hours). (B) Initial HA screen of Atlantic salmon erythrocyte elution following agglutination with 13 historical isolates of European (EU) and North American (NA) genotype. (C) Lowest haemagglutinating titre (HAU/well) at which elution was observed for erythrocytes agglutinated with elution-permissive EU isolates (EU-7-10) and NA isolates (NA-1-3). **p<0.01, ***p<0.001, ****p<0.0001, determined using multiple Mann-Whitney U tests with correction by the two-stage step-up method of Benjamini, Krieger, and Yekutieli. (D) Parallel haemagglutination reactions using cell-derived antigen and concentrated virus supernatants. White bars indicate reactions with no detectable elution (minimum 16 HAU/well), whereas red bars indicate reactions where elution was observed.

### Elution-permissive isolates show similarities in HE amino acid sequence

Phylogenetic analysis of ISAV segment 6, based on nucleotides 118-998 of the HE open reading frame, confirmed that NA isolates and EU isolates formed distinct clusters (Fig. 2A). However, the four elution-permissive EU isolates did not cluster together. In addition, elution phenotypes were not clearly associated with the position or length of the pathotypic HPR deletions (Fig. 2B-C, Table 1).

**Figure 2:**
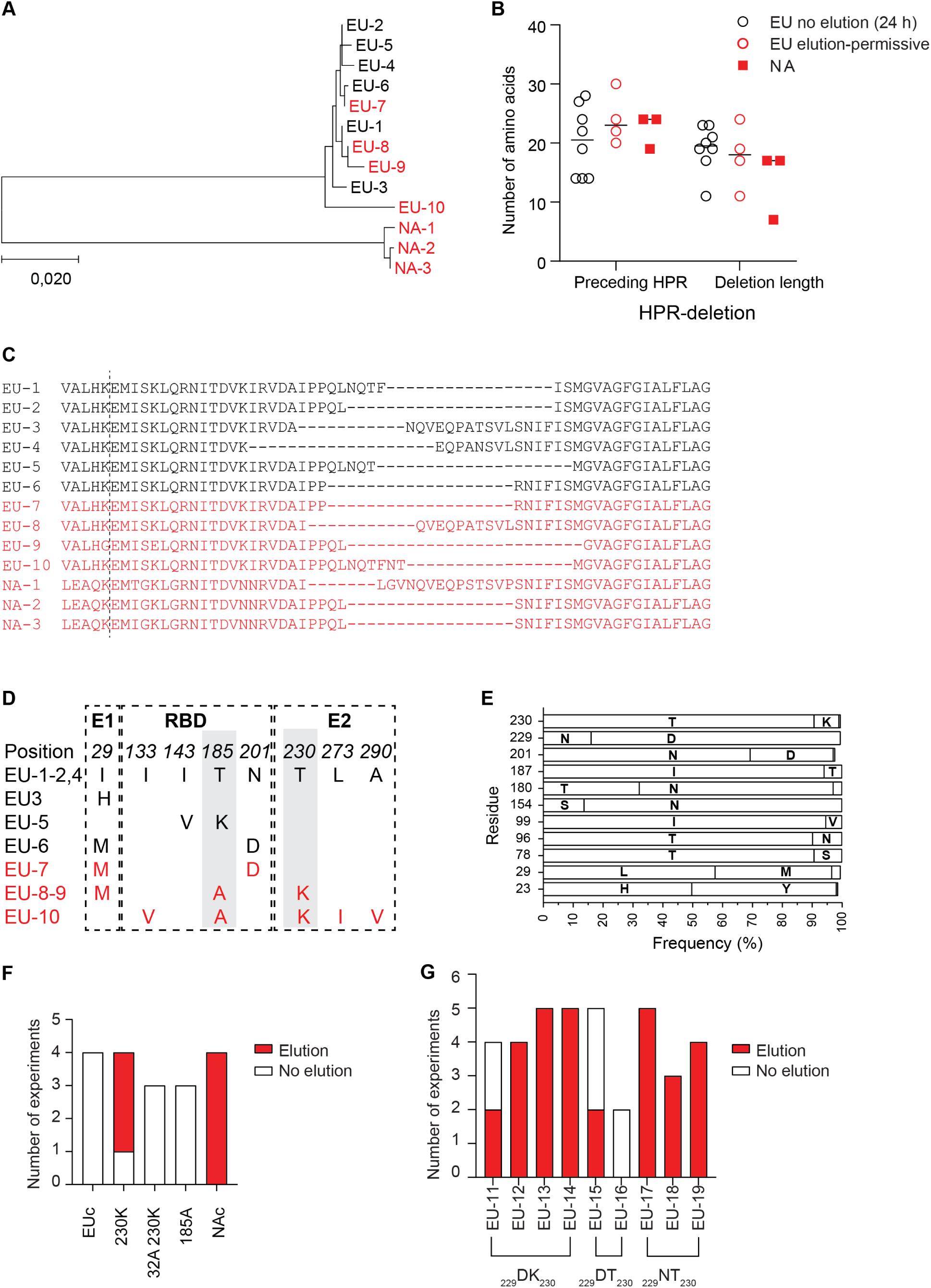
Elution-permissive ISAV isolates show common variation in HE sequences. (A) Phylogenetic clustering of ISAV isolates included in the elution screen, based on nucleotides 118-998 of segment 6. Elution permissive isolates are shown in red. The tree was constructed using the neighbour-joining method implemented in MEGA-X (version 10.1.8). (B) Properties of the highly polymorphic region (HPR)-deletion in included isolates. The number of amino acids in the HPR preceding the deletion (from and including residue 325) and deletion length (amino acids) of each isolate is shown as an individual data point. Lines indicate medians. (C) HPR alignment of the isolates included in the screen, starting from residue 320 and including the deletion site. The start of the HPR is marked by a stippled vertical line. (D) Amino acid variation detected in the receptor binding domain (RBD) and the esterase domains (E1, E2) of included European (EU) genotype isolates. HE sequences covering these regions (amino acids 17-299) from all included isolates are presented in Table S1. (E) The barplot shows HE residues with <95% conservation among ISAV sequences deposited to NCBI Genbank by the Norwegian Veterinary Institute in the period 2018-2025. Amino acid variants with >5% frequency are indicated by bold letters on the bars. (F) Elution following erythrocyte agglutination by membrane fractions of CHSE cells expressing recombinant HE, including the EU reference sequence (EUc), EUc T230K (230K), esterase-silenced EUc S32A T230K (32A 230K), EUc T185A (185A), and a North American genotype sequence (NAc). (G) Elution-permissiveness of nine recent Norwegian ISAV isolates (all EU genotype) in relation to amino acid identity at positions 229 and 230. Elution-negative isolates (E) and reactions (G) are shown in white, and elution-permissive isolates and reactions in red.

We further examined the amino acid sequence of the esterase (positions 17-126 and 213-299) and receptor-binding (positions 127-212) domains (Fig. 2, Table 1, Table S1). Three out of six EU isolates that never permitted elution shared identical amino acids across all analysed positions (EU-1, EU-2, and EU-4). This sequence was used as reference (EUc) for comparing the EU genotype isolates in our study. EU-3 diverged from the EUc sequence at a single position (Y23H), whereas EU-5 diverged at two positions (I143V, T185K). Notably, the HE sequences of EU-6, which did not permit elution, and EU-7, which permitted elution in one of four experiments, were identical. This sequence differed from the reference sequence at two positions (I29M, N210D). The I29M substitution was also present in EU-8 and EU-9; however, both isolates carried two further substitutions, T185A and T230K. The final elution-permissive isolate, EU-10, had three additional substitutions (I133V, L273I, A290V) on top of the T185A/T230K background (Table 1, Fig. 2D). To assess natural variation among HE sequences, we retrieved ISAV segment 6 sequences submitted to GenBank by the Norwegian Veterinary Institute between 2018 and 2025, all European genotype (n=205, accession numbers provided in Table S2). The highest variation in amino acid identity across the esterase and receptor-binding domains was observed at residues 23, 29, 180, 201, 229, 154, 96, 78, 230, 187, and 99, in that order (Fig. 2E). Of the variants observed among elution-permissive isolates in our screen, 29M was observed in 39%, 201D in 27.8%, and 230K in 8.3% of sequences. In addition, 229N, positioned next to 230 and present in all NA genotype isolates in our screen, occurred at 16.1% frequency (Fig. 2E). For comparison, T185A occurred in 2.9% of sequences (data not shown).

### HE 230K is sufficient to permit esterase-mediated elution

When erythrocytes were agglutinated with recombinant HE, expressed in CHSE cells, HE with the EUc sequence or a T185A substitution did not permit erythrocyte elution (Fig. 2F). By contrast, introduction of the T230K substitution permitted elution. Elution required esterase activity, as a catalytically inactive S32A/T230K double mutant did not permit elution (Fig. 2F). Consistent with results using antigen and concentrated virus, HE of the NA genotype always permitted elution (Fig. 2F).

We next tested the elution-permissiveness of an independent panel of Norwegian ISAV isolates featuring either 230K (n=4, EU-11-14), the NA genotype-associated 229N (n=3, EU-17-19), or neither of these substitutions (n=2, EU-15-16) (Fig. 2G, Table 2). All 229N isolates permitted elution in 100% of experiments (3/3, 4/4, and 5/5, respectively). Three of four 230K isolates also consistently permitted elution (4/4, 5/5, and 5/5), whereas one 230K isolate permitted elution in only 50% (2/4) of experiments. The two isolates lacking substitutions at position 229-230 diverged from the EUc sequence by three amino acids each (Table 2) and permitted elution infrequently (2/5) or not at all (0/2).

**Table 2:**
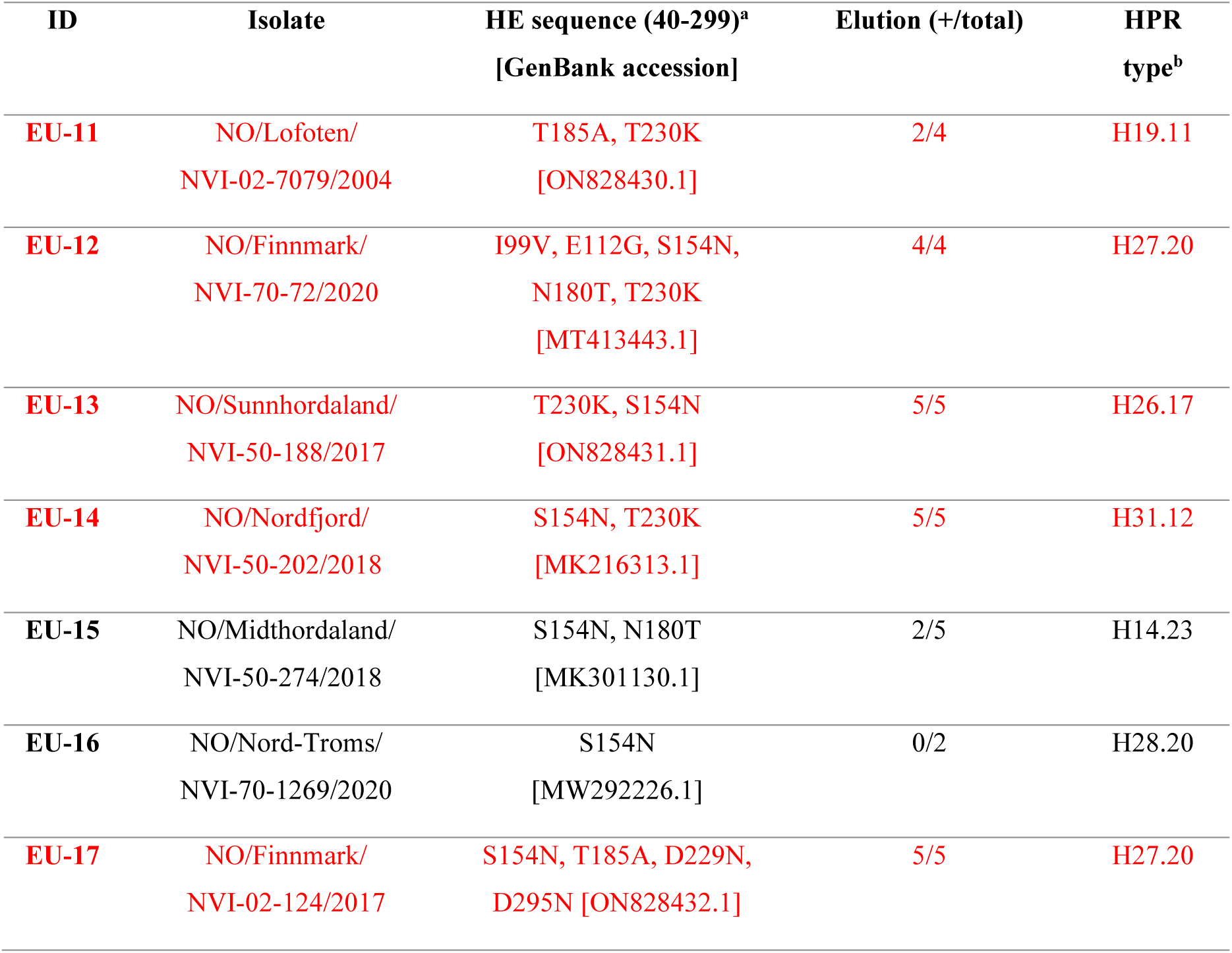
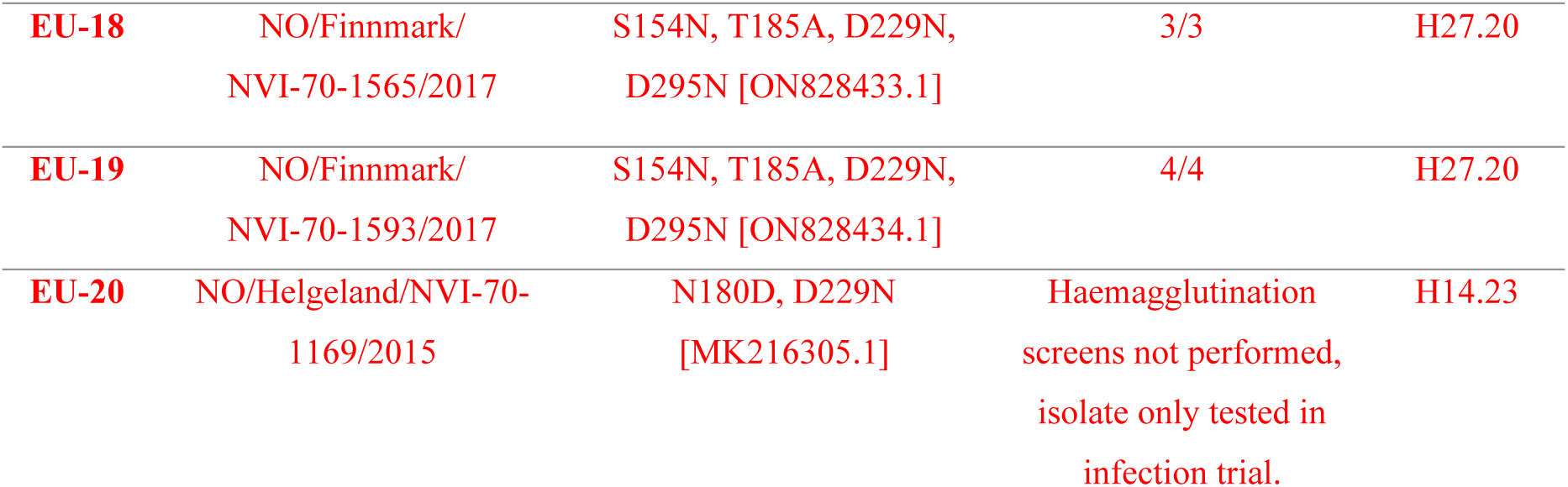
Recent Norwegian ISAV isolates included in the study. ^a^Variation from the sequence designated historical EU reference. Isolates carrying elution-permissive residues in position 229 or 230 are shown in red. ^b^HPR typing was performed as recently recommended by Gagné et al. (**22**).

### Elution patterns mirror *in vitro* and *in vivo* ISAV dissociation from Atlantic salmon erythrocytes

We next evaluated whether elution patterns observed after haemagglutination mirrored the patterns of virus – erythrocyte dissociation. We exposed erythrocytes to the same infectious dose of two ISAV isolates. Antigen from one of these isolates never permitted elution in the preceding screen (EU-5), whereas antigen from the other always permitted elution (NA-1). Erythrocyte-associated ISAV was measured by flow cytometry after 3 hours, 24 hours, 48 hours, and 72 hours. The two isolates showed clearly different patterns of erythrocyte association. While EU-5 binding increased gradually over time, NA-1 binding peaked early and subsequently declined. Increasing the EU-5 inoculum consistently increased the proportion of ISAV-bound erythrocytes but did not alter binding kinetics (Fig. 3A, Table S3). Manual inspection of flow histograms revealed tall, narrow peaks at all time points for EU-5, indicating that all erythrocytes carried comparable amounts of virus (Fig. 3B). In contrast, NA-1 produced a similarly narrow peak at 3 hours, but generated broader, flattened peaks from 24 hours onwards. This pattern was consistent with increased heterogeneity in erythrocyte-associated viral load (Fig. 3C).

**Figure 3:**
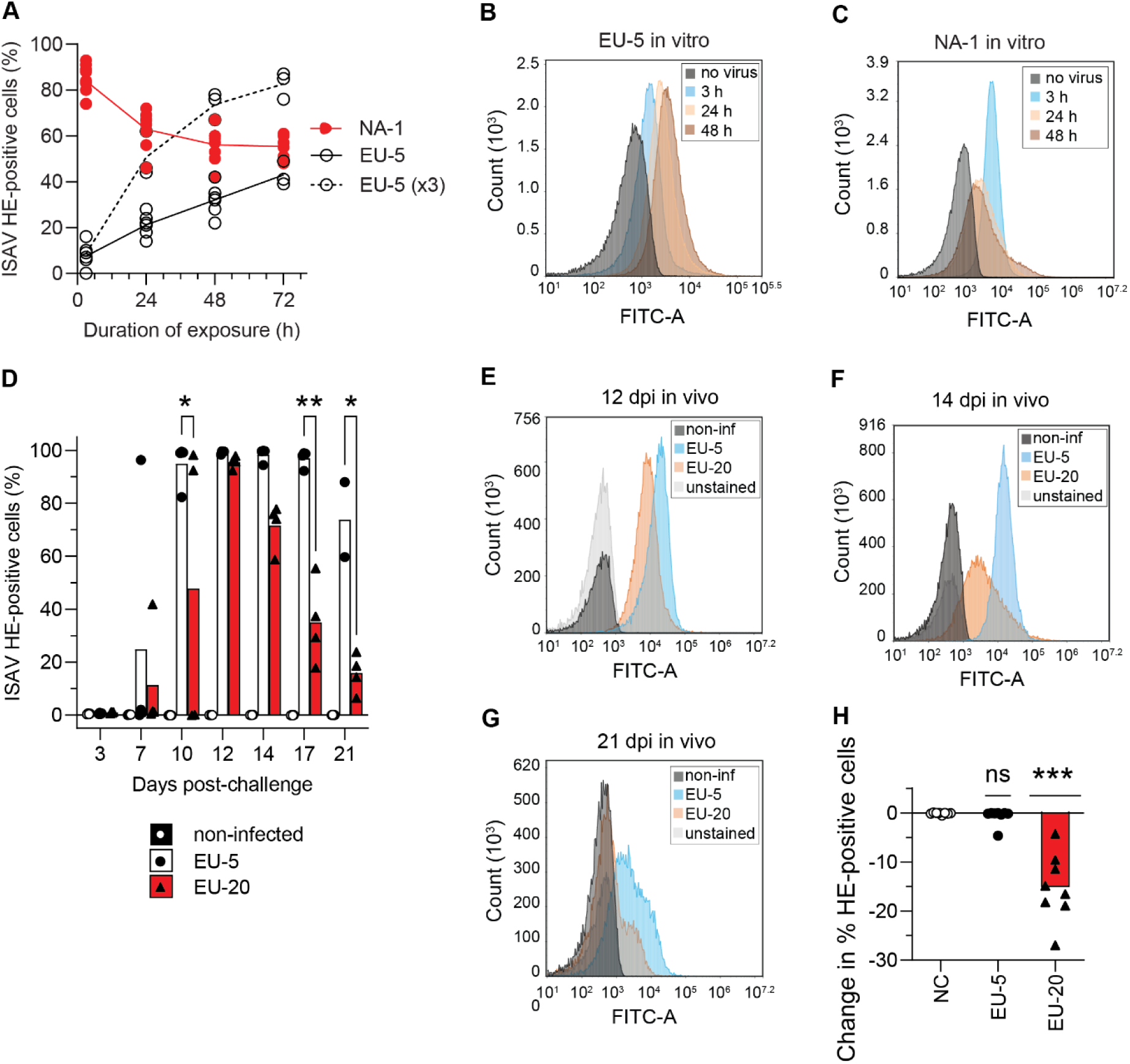
Flow cytometric detection of ISAV on Atlantic salmon erythrocytes after staining for ISAV HE. Cells were stained for ISAV HE. (A-C) Density gradient-purified Atlantic salmon erythrocytes were exposed to EU-5 or NA-1 ISAV (10^5.37^ TCID_50_/10^7^ cell, x3: 10^5.85^ TCID_50_/10^7^ cells) and stained for ISAV HE after 3, 24, and 48 hours, using mouse IgG1 (clone 3H6F8) or PBS (unstained) and goat anti-mouse IgG1-Alexa488. (A) Proportion of ISAV HE-positive cells across time points. Lines connect means. Dots show values for individual donors (n=3-9 per group). (B-C) Representative flow histograms showing the distribution of cells with different signal intensities across time points in erythrocytes from the same donor after exposure to EU-5 (B) or EU-20 (C). (D-H) Atlantic salmon blood was collected at 3, 7, 10, 12, 14, 17, and 21 days post infection (dpi) with EU-5 or EU-20. Pelleted blood cells were washed and stained for ISAV HE as described above. (D) The proportion of ISAV-positive erythrocytes in EU-5 and EU-20-infected fish across sampling points. *p<0.05, **p<0.01, calculated by two-way ANOVA with Sidak’s multiple comparisons test. Panel D data were included in a descriptive study (reference 21); mechanistic analysis is novel to this work. (E-G) Representative flow histograms showing ISAV-staining of blood cells from one non-infected, one EU-5 infected, and one EU-20 infected fish at 12 dpi (E), 14 dpi (F), and 21 dpi (G). (H) Quantification of the difference in ISAV-staining before and after shaker-incubation of washed blood cell pellets collected 12 dpi and 14 dpi (n=8 fish individuals per group, including both sampling points). ***p<0.001, calculated by one sample t test (value ≠ 0).

We next revisited data from a recent infection trial where Atlantic salmon were infected with a panel of European genotype ISAV isolates. The panel included EU-5 and EU-20, a 2015 Norwegian ISAV isolate featuring the elution-permissive HE 229N (Table 2). Both viruses caused severe infectious salmon anaemia after bath challenge with cumulative mortalities of 98% and 90%, respectively (23). Although ISAV RNA levels in blood were not significantly different (23), the proportion of ISAV-bound erythrocytes was significantly lower for EU-20 than EU-5 at 10, 17, and 21 dpi (Fig. 3D, Table S3). Mirroring observations of *in vitro*-exposed erythrocytes, flow histograms showed tall, narrow peaks in the early stages of infection (Fig. 3E-F). The proportion of ISAV-bound erythrocytes started to decline between 12 dpi and 14 dpi for EU-20, and between 17 dpi and 21 dpi for EU-5 (Fig. 3D). For both isolates, this decline coincided with a change in erythrocyte signal distribution. At 14 dpi onwards for EU-20 and 21 dpi for EU-5, the flow histogram peaks appeared broad and flattened, indicating greater heterogeneity in erythrocyte ISAV association (Fig. 3F-G). To monitor ISAV dissociation in blood from the same individual over time, blood collected 12 dpi and 14 dpi was incubated on a shaker for 24 to 48 hours before repeating flow-cytometric quantitation of ISAV-association. During the incubation period, the proportion of ISAV HE-positive cells did not decline in the samples from EU-5-infected fish. By contrast, samples from EU-20-infected fish showed an average decline of 15% (95% CI: 9-21%, p<0.001, one sample t test) (Fig. 3H).

### Erythrocyte *O*-acetylated sialic acids in Atlantic salmon, rainbow trout, and brown trout

To investigate the host determinants of ISAV-erythrocyte interactions, we analysed acetic acid-released *O*-acetylated sialic acids from salmonid erythrocytes using liquid chromatography-high resolution mass spectrometry (LC-HRMS/MS). Based on previous reports describing species-specific differences in erythrocyte interactions with the Glesvaer/2/90 ISAV isolate (EU-5) (3), we included erythrocytes from Atlantic salmon (n=7), rainbow trout (n=6) and brown trout (n=4). Although clinical disease is largely restricted to Atlantic salmon, all three species are considered susceptible to ISAV infection (24).

An overview of sialic acid derivatives contributing more than 0.5% of the total measured area in at least one individual is provided in Table 3. Representative extracted ion chromatograms are shown in Fig. 4A with measurements across samples provided in Table S4. *N*-acetylneuraminic acid (Neu5Ac) predominated in erythrocytes across all three species, accounting for more than 99.6% of the total signal area (Fig. 4B). Minor components consisted primarily of ketodeoxynonulosonic acid (Kdn) in Atlantic salmon and *N*-glycolylneuraminic acid (Neu5Gc) in rainbow trout, with only trace levels detected in brown trout (data not shown). Neu5Ac *O*-acetylation was detected in erythrocytes from all three species. Rainbow trout erythrocytes contained only mono-*O*-acetylated Neu5Ac ([M+H]^+^; *m/z* 468.1613), while Atlantic salmon erythrocytes also contained di-*O*-acetylated Neu5Ac ([M+H]^+^; *m/z* 510.1718). Di-*O*-acetylated Neu5Ac was detected in brown trout, but at much lower proportions (range: 0.15-2.7%) than Atlantic salmon (range: 7.4-19.9%). *O*-acetylated Neu5Ac peaks were identified by retention times and fragmentation patterns (Fig. S1) as compared to available reference standards, in-house reference material, and previous studies (25). The proportion of mono-4-*O*-acetylated Neu5Ac (Neu4,5Ac_2_) was similar in Atlantic salmon (range: 3.2-7.6%) and rainbow trout (range: 2.5-7.7%) erythrocytes (Fig. 4C). Consistent with reports that ISAV fail to agglutinate brown trout erythrocytes, Neu4,5Ac_2_ was undetectable in erythrocytes from three out of four brown trout and present only at trace levels (<0.3%) in the fourth individual (Fig. 4C). The proportion of mono-9-*O*-acetylated Neu5Ac

**Table 3.** Molecular formula, retention times (RT) and theoretical m/z of the protonated molecular ions [M + H]^+^ of sialic acid derivatives detected and exceeding >0.5% of measured signal in any one individual.

| Compound |  | Reference panel<sup>a,b</sup> | Formula |  | Theoretical Mass | RT |
| --- | --- | --- | --- | --- | --- | --- |
| | | | *Sia*<sup>c</sup> | *DMB-Sia*<sup>d</sup> | $[M+H]^+$<br>m/z | |
| N-Acetylneuraminic acid | Neu5Ac | a | C<sub>11</sub>H<sub>19</sub>NO<sub>9</sub> | C<sub>18</sub>H<sub>23</sub>N<sub>3</sub>O<sub>9</sub> | 426.1507 | 3.8 |
| 7-O-Acetyl-N-acetylneuraminic acid | Neu5,7Ac<sub>2</sub> | a | C<sub>13</sub>H<sub>21</sub>NO<sub>10</sub> | C<sub>20</sub>H<sub>25</sub>N<sub>3</sub>O<sub>10</sub> | 468.1613 | 4.3 |
| 8-O-Acetyl-N-acetylneuraminic acid* | Neu5,8Ac<sub>2</sub> | - | C<sub>13</sub>H<sub>21</sub>NO<sub>10</sub> | C<sub>20</sub>H<sub>25</sub>N<sub>3</sub>O<sub>10</sub> | 468.1613 | 4.8 |
| 9-O-Acetyl-N-acetylneuraminic acid | Neu5,9Ac<sub>2</sub> | a | C<sub>13</sub>H<sub>21</sub>NO<sub>10</sub> | C<sub>20</sub>H<sub>25</sub>N<sub>3</sub>O<sub>10</sub> | 468.1613 | 5.4 |
| 4-O-Acetyl-N-acetylneuraminic acid | Neu4,5Ac<sub>2</sub> | b | C<sub>13</sub>H<sub>21</sub>NO<sub>10</sub> | C<sub>20</sub>H<sub>25</sub>N<sub>3</sub>O<sub>10</sub> | 468.1613 | 5.8 |
| 7(8),9-Di-O-acetyl-N-acetylneuraminic acid | Neu5,7(8),9Ac<sub>3</sub> | a | C<sub>15</sub>H<sub>23</sub>NO<sub>11</sub> | C<sub>22</sub>H<sub>27</sub>N<sub>3</sub>O<sub>11</sub> | 510.1718 | 6.3 |
| 4,9-Di-O-acetyl-N-acetylneuraminic acid* | Neu4,5,9Ac<sub>3</sub> | - | C<sub>15</sub>H<sub>23</sub>NO<sub>11</sub> | C<sub>22</sub>H<sub>27</sub>N<sub>3</sub>O<sub>11</sub> | 510.1718 | 8.4 |

<sup>a</sup>AdvanceBio Sialic Acid reference Panel (Agilent Technologies); <sup>b</sup>Neu4,5Ac<sub>2</sub> standard (Ludger); <sup>c</sup>Before derivatization; <sup>d</sup>After derivatization; \*Based on identification by Zhang et al. (25); \*\*Mass accuracy <5 ppm

**Fig. 4:**
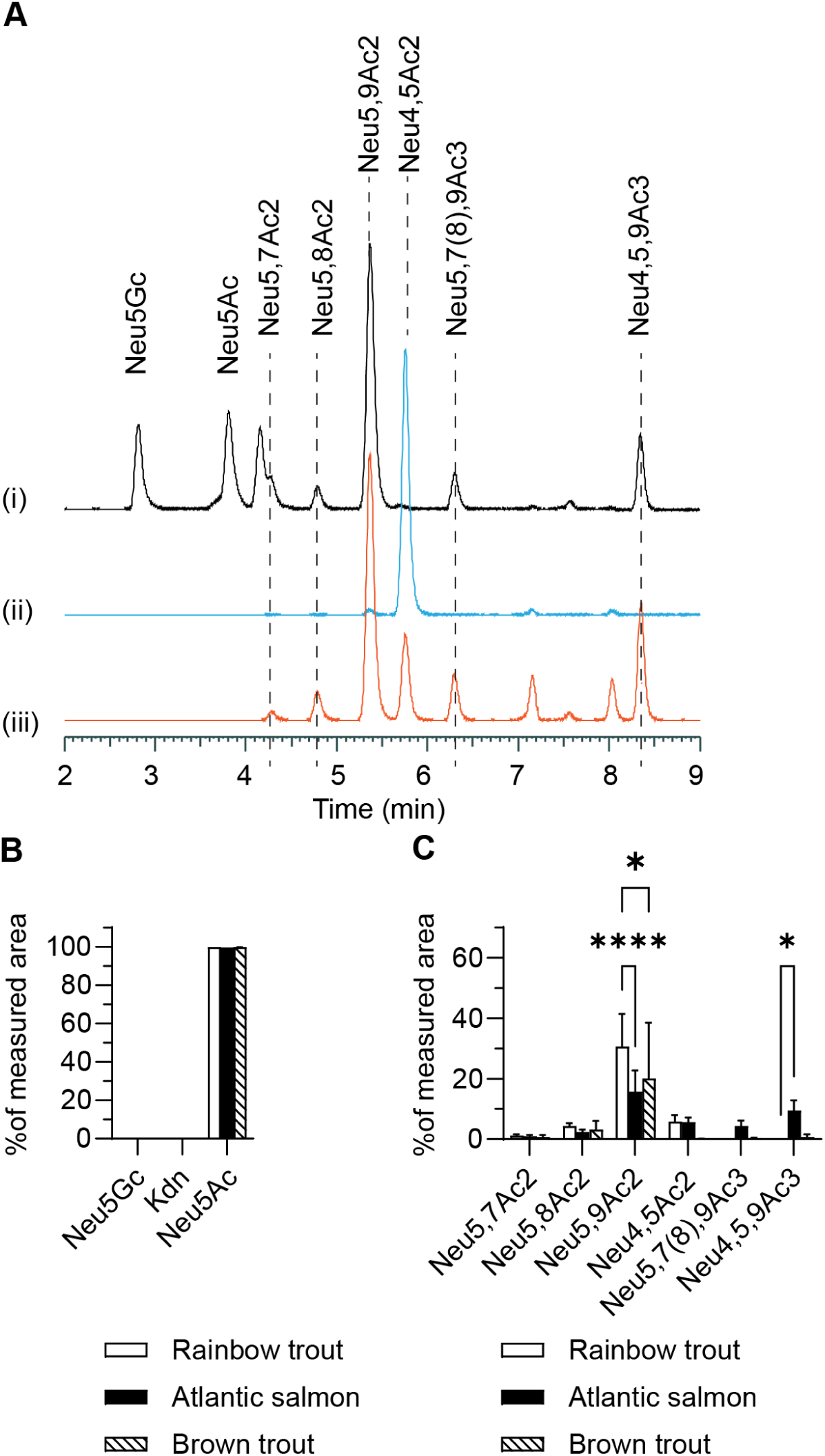
High resolution mass spectrometric analysis of salmonid erythrocyte sialic acid variants. Acetic-acid released sialic acids were extracted and labelled with 1,2-diamino-4,5-methylenedioxybenzene (DMB), before analysis on a Q Exactive™ hybrid quadrupole-Orbitrap mass spectrometer equipped with a heated electrospray ion source and coupled to a Vanquish UHPLC system. Sialic acid species were identified by comparing retention times and targeted LC-HRMS/MS fragment data (Supplementary Figure S2) with those of the available reference standards and literature. The maximum allowable mass accuracy shift between theoretical and observed masses from the full-scan LC-HRMS spectra was restricted to 5 ppm (Table 3). (A) Extracted ion chromatograms ([M+H]^+^) of sialic acids in samples from (i) sialic acid reference panel (Agilent) m/z 442.1456, 426.1507, 468.1613, 484.1562, and 510.1718; (ii) horse serum, verified by comparison to Neu4,5Ac_2_ standard (Ludger; m/z 468.1613 and 510.1718) in separate study; and (iii) Atlantic salmon erythrocyte O-acetylated Neu5Ac species m/z 468.1613 and 510.1718. (B-C) Barplots show the means and standard deviations of (B) sialic acid core structures, (C) O-acetylated Neu5Ac species in rainbow trout (n=6), Atlantic salmon (n=7), and brown trout (n=4) erythrocytes. *p>0.05, ****p<0.0001, 2-way ANOVA with Šídák’s multiple comparisons test.

was higher in rainbow trout than the two other species (Fig. 4C). Inter-individual variation was greatest for Neu5,9Ac_2_ and lowest for Neu4,5Ac_2_ in all three salmonid species.

### Loss of erythrocyte Neu4,5Ac_2_ occurs in parallel with ISAV attachment, with a delay of several days before the proportion of ISAV-bound erythrocytes declines

Finally, we extracted and analysed sialic acids from frozen blood cell pellets from the previously described trial (23). Samples included blood collected from non-infected, EU-5 infected, and EU-20 infected fish at 1 dpi, 7 dpi, and 12 dpi. The proportion of Neu4,5Ac_2_ was comparable across groups 1 dpi, but reduced in infected fish compared to non-infected fish 7 dpi and 12 dpi (Fig. 5A). Comparing EU-5 and EU-20-infected fish, Neu4,5Ac_2_ was reduced in EU-20-infected fish 7 dpi, but comparable at 12 dpi (Fig. 5A). Unexpectedly, non-infected fish showed a dramatic shift in Neu5Ac *O*-acetylation over time, increasing at the fourth and declining at the ninth carbon (Fig. 5A-B). The proportion of Neu4,9Ac_2_ was similar across groups at all time points (Fig. 5B). Regulation of Neu4,5,9Ac_3_ appeared to be more complex, with heterogeneous expression in EU-20-infected fish 1 dpi, a sharp decline mirroring the pattern of Neu4,9Ac_2_ 7 dpi, and a partial revival 12 dpi, with a non-significant trend towards being less pronounced in EU-20-infected fish (Fig. 5C). Measurements across samples are provided in Table S4. Our data reveal that infection with elution-resistant as well as elution-permissive ISAV is associated with a dramatic reduction in erythrocyte Neu4,5Ac_2_, aligning with the loss of erythrocyte ISAV binding sites observed in another trial (9). The loss of Neu4,5Ac_2_ in this trial correlated with the previously described progression of viraemia with increasing proportions of ISAV-bound erythrocytes (23) (outlined in Fig. 5D). The loss of Neu4,5Ac_2_ was almost complete by the time ISAV-binding reached its peak (Fig. 5D). Importantly, ISAV was detectable on erythrocyte surfaces for several days after the loss of Neu4,5Ac_2_ was evident. The exact duration of the delay cannot be quantified from our data, due to limited sampling points, but is estimated to be in the range from 1 to 7 days for EU-20 (7-12 dpi to 13-14 dpi) and from 5 to 13 days for EU-5 (8-12 dpi to 17-21 dpi).

**Figure 5:**
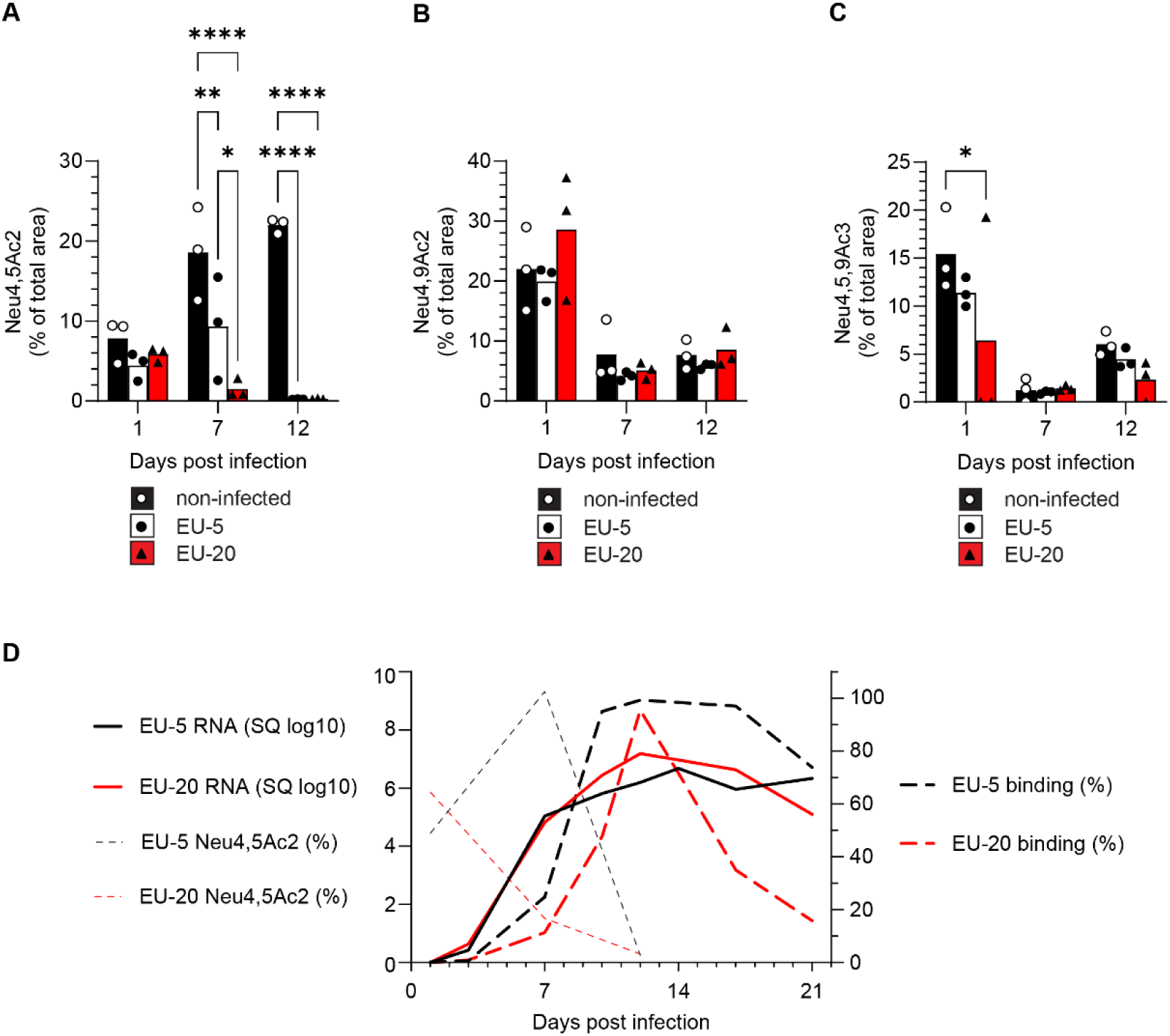
High resolution mass spectrometric analysis of erythrocyte O-acetylated sialic acids in ISAV-infected Atlantic salmon. Acetic-acid released sialic acids were extracted and labelled with 1,2-diamino-4,5-methylenedioxybenzene (DMB), before analysis a Q Exactive™ hybrid quadrupole-Orbitrap mass spectrometer equipped with a heated electrospray ion source and coupled to a Vanquish UHPLC system. Sialic acid species were identified by comparing retention times and targeted LC-HRMS/MS fragment data (Supplementary Figure S1) with those of the available reference standards and literature. Additionally, the maximum allowable mass accuracy shift between theoretical and observed masses from the full-scan LC-HRMS spectra was restricted to 5 ppm (Table 3). (A-C) Barplots showing the mean proportion of (A) Neu4,5Ac_2_, (B) Neu4,9Ac_2_, and (C) Neu4,5,9Ac_3_ in erythrocytes from non-infected, EU-5-infected, and EU-20-infected fish (n=3 per group, all included in the analysis shown in Fig. 3D), with symbols showing individual values. *p<0.05, **p>0.01, ****p<0.0001, using 2-way ANOVA with Tukey’s multiple comparisons test. (D) Timeline showing detection of ISAV RNA in blood (RNA), the proportion of ISAV-positive erythrocytes (binding), and erythrocyte Neu4,5Ac_2_ expression (Neu4,5Ac_2_) in Atlantic salmon infected with EU-5 (black) and EU-20 (red). RNA and binding data in panel D originates from a descriptive study (reference 21).

## Discussion

Our study addresses a longstanding puzzle: why Atlantic salmon erythrocytes do not elute after agglutination with the ISAV type strain Glesvaer/2/90 (EU-5 in this paper), despite the well-documented esterase activity of this virus (3, 6, 9, 10, 17). The ISAV esterase destroys its own binding sites on target cells and causes homologous attachment interference (10). Beyond this activity, the biological consequences of ISAV-mediated receptor destruction remain poorly understood. For comparison, the influenza virus neuraminidase facilitates both early and late stages of infection by promoting efficient access, entry, and release of new viral particles (11–13, 16). Here, we examine interactions between ISAV and salmonid erythrocytes to define viral and host determinants of receptor binding and disengagement. We discovered viral strain-dependent differences in elution permissiveness and erythrocyte dissociation, identifying two adjacent HE residues that accounted for most of this variability. We also observed that erythrocyte interactions with ISAV were associated with the presence of erythrocyte Neu4,5Ac_2_, both across susceptible salmonid species and during ISAV infection.

Aligning with previous work, erythrocytes agglutinated with Glesvaer/2/90 or other isolates with comparable HE sequence did not elute during 24 hours of incubation. In contrast, erythrocytes agglutinated with North American genotype isolates and a subset of European isolates did elute. Permissiveness to elution was strongly associated with amino acid identity at positions 229 and/or 230 of the ISAV HE and required a functional esterase catalytic triad. Residues 229 and 230 form part of a loop on the distal rim of the P1 pocket of the ISAV esterase, positioned away from the catalytic triad. The P1 pocket is assumed to accommodate the 4-*O*-acetyl group (5). The elution permissive T230K substitution introduces a positive charge at the rim of the pocket, whereas D229N causes loss of negative charge. Both substitutions are therefore predicted to alter electrostatic interactions with the negatively charged neuraminic acid during substrate docking, but structural studies are required to test this hypothesis. Although occasional elution was observed following agglutination by ISAV antigen with HE changes at other positions, these effects were inconsistent and of lesser magnitude.

The evidence that Neu4,5Ac_2_ serves as an initial attachment receptor for ISAV on erythrocytes is strong. First, the dynamics of erythrocyte Neu4,5Ac_2_ elimination in our study closely mirror the previously described loss of ISAV binding sites at erythrocyte and vascular surfaces during infection (9, 10). Moreover, the distribution of Neu4,5Ac_2_ in salmonid erythrocytes correlated with their ability to be agglutinated by ISAV (3). High-resolution mass spectrometric analysis showed that rainbow trout and Atlantic salmon erythrocytes, both reported to be agglutinated by ISAV, expressed comparable levels of Neu4,5Ac_2_. In contrast, brown trout erythrocytes, which do not support ISAV agglutination, expressed little or no Neu4,5Ac_2_. Notably, erythrocytes primarily act as reservoirs for circulating virus particles and only occasionally express ISAV RNA and protein, whereas replication in endothelial cells is essential for the progression of disease (26–28). Further studies should therefore determine whether Neu4,5Ac_2_ also mediates attachment to the vascular endothelial surface.

Endothelial-produced ISAV particles are released into the circulation, causing progressive viraemia with extensive erythrocyte binding. The proportion of circulating Atlantic salmon erythrocytes bound by ISAV declines with time during experimental infection, even for elution-resistant strains (9, 23). Here, we confirm this observation and show that the decline was associated with increased heterogeneity in erythrocyte ISAV signal. This suggest that the decline cannot be explained entirely by removal of ISAV-bound erythrocytes from the circulation (26), but also reflects a reduced capacity of circulating erythrocytes to bind the virus. The time window for ISAV detection on circulating erythrocytes was significantly narrower for a strain carrying an elution permissive HE variant than for Glesvaer/2/90. Loss of erythrocyte Neu4,5Ac_2_ occurred in parallel with initial ISAV binding. Moreover, a delay was observed between the loss of Neu4,5Ac_2_ and the subsequent decline in ISAV-bound erythrocytes for both strains, although this delay was most pronounced for Glesvaer/2/90. Together, these observations indicate that ISAV receptor destruction is an early event in viraemia but do not explain the persistence of ISAV on erythrocyte surfaces after Neu4,5Ac_2_ hydrolysis.

While our cell-based approach does not provide evidence for the mechanism underlying this persistence, the findings nevertheless support the generation of informed hypotheses. Given the ∼29 Å distance between the ISAV receptor-binding domain and the esterase pocket (5), it appears unlikely that the esterase can hydrolyse Neu4,5Ac_2_ molecules that are directly engaged by the receptor-binding domain. It is therefore likely that, following receptor engagement, the esterase hydrolyses Neu4,5Ac_2_ in the vicinity of the bound receptor. ISAV may then remain bound until viral entry occurs, or receptor disengagement takes place. Following disengagement, ISAV may either dissociate or re-engage, depending on the local availability of Neu4,5Ac_2_ on the cell surface. Slow receptor disengagement could be one explanation for the observed delay between receptor loss and the decline in ISAV-bound erythrocytes. An alternative, non-exclusive explanation is that initial engagement with Neu4,5Ac_2_ facilitates binding to a secondary attachment factor that is less susceptible to ISAV esterase activity. In this context, it would be of interest to determine whether Neu4,5,9Ac_3_ supports ISAV attachment and whether it is susceptible to hydrolysis by the ISAV esterase.

The initial discovery that Atlantic salmon erythrocytes did not elute from ISAV agglutination was based on EU-5 (3). Following this report, unpublished observations confirmed the elution-resistant phenotype in a range of ISAV isolates from severe field outbreaks in the 1990s. We show that both elution resistant and elution permissive ISAV strains can cause severe disease with high mortality in experimental settings. However, the trial had limited power to resolve the role of the ISAV esterase in disease outcome and transmission, because of co-existing genomic differences between the viral strains (23). Accordingly, further studies are required to determine whether the differences in ISAV esterase activity and erythrocyte dissociation affect viral fitness and the outcome of infection, including transmission. Although ISAV-erythrocyte interactions provide a useful system for studying receptor binding and destruction, their direct relevance to ISAV pathogenesis remains uncertain. Erythrocyte binding only becomes detectable after the replication in endothelial cells has started, suggesting a higher relevance for viral clearance than for establishing infection (9, 26). Accordingly, decoy receptors in other anatomical compartments may be more important for the early stages of infection that determine outcome. Neuraminidase activity is critical for influenza A virus mucus penetration, with higher activity correlating with lesser sensitivity to mucus inhibition (16). Given the importance of mucus barriers for early infection and transmission in other viral infections, a similar role for the ISAV esterase warrants investigation.

Taken together, our study addresses the longstanding conundrum of why Atlantic salmon erythrocytes do not elute following ISAV-agglutination and reveals functional strain differences with a defined genetic basis, occurring in 25% of current Norwegian ISAV strains. Moreover, our analysis of erythrocyte sialic acids across ISAV-susceptible salmonid species and their association with ISAV-binding underscores the crucial role of Neu4,5Ac_2_ as an ISAV attachment receptor. At the same time, our findings suggest that additional host molecules or binding modes contribute to prolonged ISAV persistence on erythrocyte surfaces after initial attachment. By generating new mechanistic insights into how ISAV attaches to and is released from host cells, our study provides a framework for dissecting ISAV receptor usage and its contribution to host susceptibility.

## Materials and methods

### Viruses

ISAV strains were isolated from farmed Atlantic salmon as part of outbreak investigations. Propagation of viruses for experiments were done in Atlantic salmon kidney (ASK) cells (29) infected at approximately 80% confluence. ASK were maintained in Leibovitz-15 medium (Lonza) supplemented with foetal bovine serum (FBS, 10%, Lonza), L-glutamine (Lonza, 4mM), and penicillin/streptomycin/amphotericin (Lonza, 1%), hereafter referred to as growth medium, in closed cap tissue culture flasks at 20 °C without exogenous supplementation of CO_2_, and split 1:3 every 14 days. After infection, cells were incubated at 15 °C. Antigen for haemagglutination-elution assays was prepared by collecting membrane fractions of infected ASK cells, as previously described (27). Cells in 75 cm^2^ tissue culture flasks were harvested when cytopathic effects were evident, but the cells had not yet detached, typically 3-7 dpi. Cells were washed and scraped on ice, and cell pellets containing cellular-expressed viral membrane glycoproteins were washed, resuspended in 0.5 mL PBS, and subjected to three rounds of freeze-thawing. Viruses were obtained by harvesting supernatants when cytopathic effects were close to complete, typically 14-28 dpi. Infective titres were determined by inoculating serial dilutions of supernatants or blood (10 µL heparinised blood was harvested into 500 µL L-15 medium and kept at -80 °C until inoculation on cells) in quadruplicate wells of ASK cells cultured in 96-well microtiter plates. Acetone-fixed cells were incubated with IgG_1_ specific to the ISAV nucleoprotein (P10, Aquatic Diagnostics Ltd, 0.4 µg/mL, 60 min, RT), washed (PBS × 3), and incubated with Alexa488-labelled goat anti-mouse IgG (A11001, Thermo Fisher Scientific, 5 µg/mL, 45 min, RT), and titres were calculated by the modified Kärber method, as previously described (3).

### Genomic sequencing and sequence analyses

Near full-length ISAV segment 6 was amplified using the SuperScript IV^TM^ One-Step RT-PCR System (Thermofisher) and subjected to Sanger sequencing to identify the segment 6 genotype as previously outlined (30, 31). Amplification was achieved using the following forward/reverse primers: 5’-AGCAAAGATGGCACGATTCATAATTT-3/5’-CGTACAACATCAAGAACGTCTTC-3’, generating a product of 1307 nt. Using the forward primer and an additional sequencing primer (5’-GTGTCAGACACCTTTGAAGTGAG-3’), the purified amplicons were sequenced using the BigDye^TM^ Terminator Cycle sequencing kit v.3.1 (Thermo Fisher Scientific) and analysed on a SeqStudio Genetic Analyzer (Thermo Fisher Scientific). The resulting sequences were analysed with Sequencing Analysis Software 6.0 (Thermo Fisher Scientific) and assembled using the CLC Main Workbench 21.0.3 (Qiagen). To determine the evolutionary relationship between historical ISAV strains, a neighbour-joining phylogenetic analysis based on nucleotides 118-998 of the segment 6 open reading frame was performed in MEGA-X (version 10.1.8).

The HE sequences of all historical isolates are presented in Table S1, while accession numbers for recent Norwegian isolates are provided in Table 2. Manual HPR typing was performed as recently recommended by Gagné et al. (**22**). Briefly, the designation begins with H (for HE), followed by the number of amino acids from, and including, residue 325 to the start of the HPR deletion, and the number of deleted amino acids, separated by a full stop. For example, EU-5 is designated HPR type H27.20, indicating 27 amino acids from residue 325 to the deletion site and a deletion of 20 amino acids.

To examine the natural variation in HE sequence, ISAV segment 6-sequences deposited by the diagnostic services at the Norwegian Veterinary Institute were retrieved from the NCBI GenBank 30.04.2026 (accession numbers provided in Table S2). Sequences were aligned and translated in Jalview (version 2.11.5.1) (32), and annotations were exported. A visual presentation of highly variable sites within the esterase and receptor-binding domains (<95% consensus, >95% occupancy in dataset) was generated in GraphPad Prism (version 10.3.1).

### Generation of recombinant HE for haemagglutination-elution assay

Codon optimised sequences of the open reading frame encoding ISAV HE of European genotype, including 32A, 185A, and 230K variants, and North American genotype, were inserted in the pcDNA3.1 (+) vector commercially, and delivered transfection-ready (GeneCust, Boynes, France). Monolayers of CHSE-214 cells were cultured until 90-100% confluent and detached by Trypsin EDTA (Lonza Bioscience). Four transfection reactions, each with 10^6^ cells and 10 µg plasmid DNA, were performed, using the Neon 100 µL Transfection System (Invitrogen, Waltham, MA, USA, three pulses at 1600 V and 10 ms width). The transfected cells were pooled and incubated 24 hours in antibiotic free medium, then another 24 hours in culture medium. Membrane fractions were collected as described above for infected cells.

### Fish and sample collection

Atlantic salmon and rainbow trout (AquaGen, Trondheim, Norway) were purchased from the aquaculture research station: Center for Sustainable Aquaculture (Norwegian University of Life Sciences [NMBU], Ås, Norway). Fish were maintained on a 24 hour light photoperiod in circular tanks in a temperature-controlled (14±1 °C) fresh water recirculatory aquaculture system and fed a standard salmon diet in excess from automatic belt feeders. Blood for *in vitro* experiments was collected in heparinised containers. Caudal vein terminal blood sampling was performed on fish anaesthetised by immersion in tricaine mesylate (100-200 mg/L). The infection trial (NMBU) has been described before (23). Briefly, Atlantic salmon (50-80 g) were immersed in ISAV-spiked water (10^3.5^ TCID_50_/mL) for 2 hours, transferred to two 500 L tanks per strain for sampling (n=70) and observation (n=40), respectively. Fish were maintained at 12 °C, 12:12 hours photoperiods, and about 80% oxygen saturation throughout the trial. Eight ISAV strains were tested in parallel, including EU-5 and EU-20. An additional tank containing 40 non-infected fish served as controls. Fish were inspected daily for clinical signs and mortality. Samples from brown trout were obtained by the Norwegian Veterinary Institute as part of diagnostic investigation at the Norwegian Forest Museum (Elverum, Norway) and routine health assessment of brood stock of anadromous brout trout (sea trout) at the live gene bank for wild salmonids (Eidfjord, Norway).

### Haemagglutination/elution testing and binding assays

For haemagglutination/elution assays, the blood was washed twice in cold PBS, resuspended 1% in PBS 0.5% BSA, and used the same day. The assays were performed in a V-bottom 96-microwell plate, prefilled with PBS (50 µL/well). Virus or antigen was added to the first column (50 µL/well), serially diluted 1:2 to column 11, with column 12 containing PBS only. Next, erythrocytes were added (1% suspension, 50 µL/well), incubated (30 min, room temperature), and haemagglutination titres were determined. Elution, defined as re-appearance of the erythrocyte button (Fig. 1A), was monitored over a period of 24 hours (incubation in the dark, room temperature). For the *in vitro* investigation of ISAV binding by flow cytometry, erythrocytes were isolated by density gradient centrifugation. Briefly, 0.5 mL blood was diluted in 5 mL phosphate-buffered isotonic saline (PBS), layered on top of 7.5 mL 51% Percoll Plus (Sigma-Aldrich), and centrifuged without breaks (500 × *g*, 20 min, 10 °C). Pellets were washed twice in cold PBS, resuspended 2 × 10^7^ cells/mL in growth medium, and maintained on a microplate shaker (150-200 rpm, 15 °C) until use. Exposure to ISAV was performed by pelleting and resuspension of 1 × 10^7^ cells in growth medium containing 10^5.37^ or 10^5.85^ TCID_50_ ISAV and incubation on a shaker for 1, 24, 48, or 72 hours before staining and flow cytometric analysis. For investigation of *ex vivo* ISAV dissociation from blood cells from infected fish, blood was sampled at 12 dpi and 14 dpi (10 µL blood per fish in 1 mL growth medium) and incubated on a microplate shaker (150-200 rpm, 15 °C) for 24-48 hours before staining and flow cytometric analysis.

### ISAV-staining and flow cytometry

After ISAV-incubation, live density-purified erythrocytes were transferred to duplicate wells of a 96-well microplate (2×10^6^ cells/well). The plate was centrifuged (500 ×*g*, 15 °C, 5 min), supernatants discarded, and the pellets were resuspended in 50 µL primary antibody (mouse IgG1 anti-ISAV HE, clone 3H6F8 (33), 1/100) or PBS. Heparinised blood from infected fish was stored at 4 °C until immunostaining and analysis and washed once by the addition of 1 mL PBS and centrifugation (500 ×*g*, 15 °C, 5 min), 10 µL pelleted cells were suspended in 1000 µL PBS, and 100 µL of this suspension was transferred to duplicate wells in a 96-well microplate. The plate was centrifuged (500 ×*g*, 15 °C, 5 min), supernatants discarded, and the pellets were resuspended in 50 µL primary antibody (mouse IgG1 anti-ISAV HE, clone 3H6F8 (33), 1/10 to enhance detection) or PBS. Subsequent steps were similar for blood pellets and purified erythrocytes. After incubation with the primary antibody (room temperature, 60 min), 100 µL PBS was added to each well before centrifugation and two additional washes, each in 150 µL PBS, resuspension with goat anti-mouse IgG1-Alexa488 (Molecular Probes, #A11001, 5 ug/mL), incubation (room temperature, shielded from light, 45 min), wash in PBS as above, resuspension in 100 µL PBS, and flow cytometric measurement of fluorescence in at least 20,000 cells (10,000 cells for PBS-stained wells) by a Novocyte Flow Cytometer (Agilent). To account for potential strain-specific differences in antibody affinity, the signal was expressed as percentage of ISAV HE-positive erythrocytes, exceeding the signal range of cells from non-infected fish.

### Liquid chromatography-high-resolution mass spectrometry (LC-HRMS)

The acetic acid-released sialic acids were analyzed using a Q Exactive™ hybrid quadrupole-Orbitrap mass spectrometer equipped with a heated electrospray ion source (HESI-II) and coupled to a Vanquish UHPLC system (Thermo Fisher Scientific). Chromatographic separation was achieved using an ACE Excel 2 C18-AR column (2 μm, 2.1 × 100 mm) maintained at 40°C. Methanol (MeOH) and water (H₂O), each containing 0.1% formic acid, were used as mobile phases A and B, respectively. The compounds of interest were separated at a flow rate of 0.3 mL/min with an injection volume of 1 μL. The elution gradient began with 20% mobile phase B and followed a programmed gradient as time (min)/B (%): 9/36, 12/50, and 12.2/90. The total run time was 16 minutes including equilibration. The HESI-II interface was operated in positive ion mode at 300°C with a spray voltage of 3.2 kV, capillary temperature of 280 °C, sheath gas flow rate of 35 L/min, auxiliary gas at 10 L/min, and S-lens RF at 55. Mass spectra were acquired in full scan mode with ddMS^2^-top5 and an inclusion list for targeted masses of interest (Table 1). Full scan settings included 70,000 FWHM resolution, AGC target of 1 × 10^6^, max IT of 256 ms, and scan range of m/z 300–600. HRMS/MS spectra were acquired at 17,500 FWHM using HCD with stepped NCEs of 20, 35, and 60 eV. AGC target, IT, and isolation window were set to 5 × 10^4^, 150 ms, and *m/z* 1.6. The instrument was calibrated once a week both in positive and negative mode using an external calibration solution (Pierce®, ThermoScientific, San Jose, CA, USA) according to the manufacturer’s recommendations. Processing of high-resolution MS data was carried out using Xcalibur 4.2.28.14 (Thermo Fisher Scientific). The identification of known sialic acids was achieved by comparing retention times and LC-HRMS/MS fragment data (Supplementary Fig. S1) with those of the available reference standards. In cases where reference materials were unavailable, potential sialic acids were identified by comparing the mass spectra with both negative and positive controls to detect the peak of interest. Additionally, the maximum allowable mass accuracy shift between theoretical and observed masses from the LC-HRMS spectra was restricted to 5 ppm (Table 1).

### Sample preparation

HRMS analysis was performed following mild acid hydrolysis and derivatisation of the released sialic acids with 1,2-diamino-4,5-methylenedioxybenzene (DMB). Briefly, 20 µL of 2M acetic acid was added to 10 µL of the sample and incubated at 80 °C for 2 hours. After hydrolysis, the samples were cooled, vortexed, and centrifuged at 10,000 x g for 3 minutes. For labelling, 5 µL of the sample was mixed with 20 µL of freshly prepared 7 mM DMB solution containing 1.3 M acetic acid and 0.74 M 2-mercaptoethanol. After a 3-hour incubation at 50°C, the reaction was terminated with 480 µL of water (H₂O). The final solution was desalted using Phenomenex Strata-X C18-E cartridges preconditioned with 1 mL MeOH and 1 mL H₂O. After washing with 5% MeOH, the sample was eluted with 1 mL of 100% MeOH. The samples were evaporated under a nitrogen stream and re-dissolved in 30% MeOH (v/v) before HRMS analysis. The reference material was supplied as a dried mixture of sialic acid derivatives, totalling approximately 1.25 nmol (AdvanceBio Sialic Acid Reference Panel, Agilent Technologies) or 1 nmol (Neu4,5Ac_2_ sialic acid standard, Ludger) per vial. Before analysis, 0.012 mL of water was added to the dried material, and the mixture was vortexed for 30 seconds. A 0.005 mL aliquot was then used for DMB labelling.

## Author contributions

Conceptualisation: JHF, KF, Data curation: JHF, PEP, MMD; Formal analysis: JHF, LI; Funding acquisition: JHF, SW, SP, DC, KF; Investigation: JHF, AMSA, LI, FBP, IAH, SW, PEP, MMD, LF, BT; Methodology: JHF, LI, SW, DC, KF, Project administration: JHF; Resources: JHF, LI, DC, AG, KF; Supervision: JHF, SP, DC, AG, KF; Validation: JHF, AMSA, LI, PEP, MMD; Visualization: JHF, LI; Writing – original draft: JHF, Writing – review and editing: All authors.

## Acknowledgements

The authors would like to thank Torfinn Moldal, Åse Helen Garseth, and Brit Tørud (Norwegian Veterinary Institute), staff at the NMBU Center for Sustainable Aquaculture, NMBU infection aquarium, the Norwegian Forest Museum, and the Gene bank for wild Atlantic salmon (Norwegian Environment Agency, Eidfjord, Norway) for kind advice and assistance. The authors would also like to acknowledge the Red Flag project funded by the Research Council of Norway (Grant ID#302551) for methodological support for erythrocyte experiments.

## Ethics statement

Sample collection from fish and the ISAV infection trial were performed according to protocols pre-approved by the Norwegian Food Safety Authority (FOTS ID#24382, ID#30432, ID#30501). The facilities for the infection trial were operated in compliance with OECD principles of Good Laboratory Practice and Guidelines to Good Manufacturing Practice issued by the European Commission.

## Data availability

All data generated and analysed within the current study are available within the manuscript and its supplementary files.

## Supplementary material

**Fig. S1:**
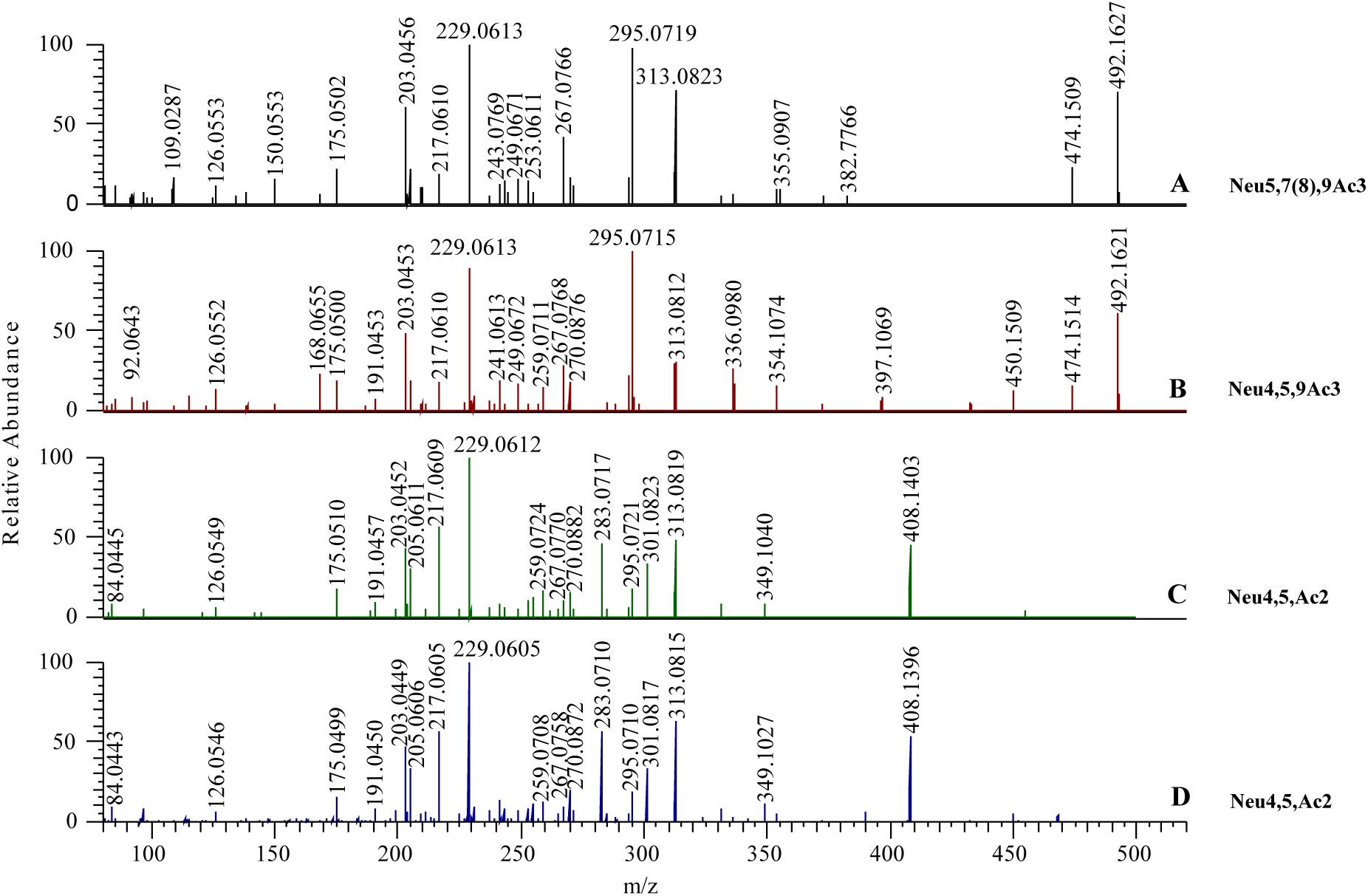
Supplementing Fig. 4. Representative images showing LC-HRMS/MS fragment data of peaks of interest. Panels represent (A) Neu5,7/8,9Ac_3_ (m/z 510.1718), with a lack of m/z 259.1 indicating a substitution in the C7 position, (B) Neu4,5,9Ac_3_ (m/z 510.1718), with the presence of m/z 259.1 indicating a lack of substitution in the C7 position and a lack or low abundance at m/z 283.07 and 301.08 indicating a substitution in the C8 or C9 position, and (C) Neu4,5Ac_2_, with a lack of m/z 450.2 [M+H-18]^+^, abundant m/z 283.07, and identified in horse serum used as internal control, as well as Neu4,5Ac_2_ standard (D; Ludger).

***Table S1****. Supplementing Fig. 2 and Table 1. Provides HE sequences of residues 17-299 (covering the esterase and receptor-binding domains) of historical ISAV isolates included in the elution screen*.

***Table S2.*** *Supplementing Fig 2E. Provides the NCBI GenBank accession numbers of ISAV segment 6 sequences used to assess the variation in HE amino acid identity in field samples from Norwegian farms spanning 2018-2025 (n=205)*.

***Table S3.*** *Supplementing Fig 3. Flow cytometric quantification of ISAV-HE across samples. Density gradient-purified Atlantic salmon erythrocytes exposed to EU-5 or NA-1 ISAV (105.37 TCID50/107 cells, x3: 105.85 TCID50/107 cells) or pelleted blood cells from infected fish were washed and stained for ISAV HE, using mouse IgG1 (clone 3H6F8) or PBS (unstained) and goat anti-mouse IgG1-Alexa488. Gating based on forward scatter (FSC) and side scatter parameters were used to identify cells and exclude debris, and doublets were excluded using FSC-A versus FSC-H gating. Fluorescence thresholds for Alexa Fluor 488 positivity were defined based on unstained and PBS-treated controls to account for autofluorescence and background signal. Fluorescence was measured for at least 20,000 cells (10,000 cells for negative control, PBS-stained wells) using a Novocyte Flow Cytometer (Agilent). To account for potential strain-specific differences in antibody affinity, the signal was expressed as the percentage of ISAV HE-positive cells exceeding the fluorescence range of cells from non-infected fish. Flow cytometry data were analysed using NoVo Express (Agilent) software, and consistent gating strategies were applied across all samples*.

***Table S4.*** *Supplementing Fig 4-5. Mass spectrometric quantification of peak areas corresponding to individual sialic acid species across samples. Acetic-acid released sialic acids were extracted and labelled with 1,2-diamino-4,5-methylenedioxybenzene (DMB), before analysis a Q Exactive™ hybrid quadrupole-Orbitrap mass spectrometer equipped with a heated electrospray ion source and coupled to a Vanquish UHPLC system. Sialic acid species were identified by comparing retention times and targeted LC-HRMS/MS fragment data (Supplementary Figure S1) with those of the available reference standards and literature. Additionally, the maximum allowable mass accuracy shift between theoretical and observed masses from the full-scan LC-HRMS spectra was restricted to 5 ppm (Table 3)*.

